# A random-matrix theory of the number sense

**DOI:** 10.1101/160895

**Authors:** T. Hannagan, A. Nieder, P. Viswanathan, S. Dehaene

## Abstract

Number sense, a spontaneous ability to process approximate numbers, has been documented in human adults, infants and newborns, and many other animals. Species as distant as monkeys and crows exhibit number-selective neuronal activity. How number sense can emerge in the absence of learning or fine tuning is currently unknown. We introduce a random matrix theory of self-organised neural states where numbers are coded by vectors of activation across multiple units, and where the vector codes for successive integers are obtained through multiplication by a fixed but random matrix. This cortical implementation of the “von Mises” algorithm explains many otherwise disconnected observations ranging from neural tuning curves in monkeys to looking times in neonates and cortical numerotopy in adults. The theory clarifies the origin of Weber-Fechner’s law and yields a novel and empirically validated prediction of multi-peak number neurons. Random matrices constitute a novel mechanism for the emergence of brain states coding for quantity.

## Main Text

What is the origin of our ability to represent numbers? The evidence is now overwhelming for an approximate number system (*1*) shared between several species and relying on number neurons distributed in parietal and prefrontal cortex *(2)*. Number neurons are cells whose firing varies systematically with the number of objects or events, typically with coarse tuning around a cell-specific preferred number, independently of low-level properties (e.g. size, spacing, intensity). Yet their tuning properties are debated *(3)*, and the precise circuit that allows them to exhibit numerical tuning is still unknown.

An outstanding theoretical question about number sense concerns its presence in humans and other animals prior to learning. Number neurons have been found in untrained animals *(4)* and numerical discrimination is present in human neonates *(5)*. Such findings suggest that self-organizing properties of neural circuits, early on during development, are responsible for the emergence of number-coding neurons. Furthermore, the number sense possesses very specific empirical properties, labelled here as D1-7, that any convincing theory should confront: (**D1**) The number sense obeys Weber-Fechner’s law: our ability to discriminate between numbers varies as the log of their ratios *(6)*. (**D2**) Number neurons are tuned to a broad range of preferred numerosities up to 30 items *(7)*, including quantity zero *(8-10)*, with log-Gaussian tuning curves *(7,11)*, and are found in monkeys even in the absence of numerical training *(4)*. (**D3**) Neurons tuned to low and high numbers are more frequent than neurons tuned to the middle of the tested range *(7, 11)*. (**D4**) Monkeys trained to order small numerosities spontaneously generalize to larger numerosities *(12, 13)*. (**D5**) Neonates, hours after birth, can discriminate numbers that are in a sufficiently large ratio *(5)*. (**D6**) In the course of development, number sense becomes increasingly precise *(14, 15)*. (**D7**) In human parietal cortex, voxels selective for similar numbers are more likely to be contiguous, forming a macroscopic cortical map [“numerotopy”, *(16)*].

Previous computational models based on number detector units have mostly focused on properties **D1-3**, but have typically failed to address the broader issue of the emergence of these properties without training. In order to obtain operational number detectors, these models relied either on fine-tuned hand-wired circuitry *(17, 18)*, or on learning through exposure to many numerical sets *(19, 20)*. An exception is *(21)*, which reported evidence for **D1** and **D2** in a randomly connected network of spiking neurons subject to short-term plasticity. Indeed, random networks are known to mimic the selectivity and sparsity of neural activations in particular regimes *(22)*.

Here we present a random-matrix theory of number sense that approaches numerosity at the level of vector states. We reasoned that, if each number is represented by a vector of activity across a large number of neurons, passing from one number to the next could be achieved by matrix multiplication (see *ref. 23* for a related approach in computational linguistics). We here consider the simplest possibility, which is that this successor matrix is random. We therefore envisaged that the approximate number system could be built on the powers of a certain type of random matrix: starting from some initial vector, the vectors coding for successive numbers would be generated by successive multiplications by the same fixed matrix (corresponding to the successor function +1, a foundational concept of Peano arithmetic). We show how all of the above properties **D1-7** inevitably follow from this simple theory. We implement the theory in two different models with different degrees of biological realism, that both possess very little initial structure and draw no statistics at all from the environment.

## Results

### The minimal number sense model

We start by presenting the minimal implementation of the theory: the minimal number sense model, illustrated in Fig. 1. The model is entirely described by a n × n connectivity matrix M, an initial activation vector S_0_ of dimension n, and discrete time dynamics for updating unit activations. M represents a random pattern of connectivity between number neurons which is stable for the timescale of an experiment. Each entry M_ij_ is a Gaussian random variable whose amplitude drops exponentially with the distance between units. The adjacency matrix of the minimal model is a well-studied object in mathematics–a random band matrix *(24)-* which has been used to describe spatially constrained interactions in several physical systems (see *ref. 25* for an exposition). In our case, such a random band matrix is intended to capture both the sparsity of functional connectivity in the nervous system, and its exponential decrease with the distance between neocortical neurons *(26)*. S_0_ represents the initial firing rates of number neurons in absence of stimulus, in line with the fact that neurons can be selective to numerosity zero *(8-10)*. We further assume that S_0_ is localized: its non-zero components are clustered together on the left side of the line (although even this assumption can be relaxed: see Supplementary Note 1).

**Fig. 1.**
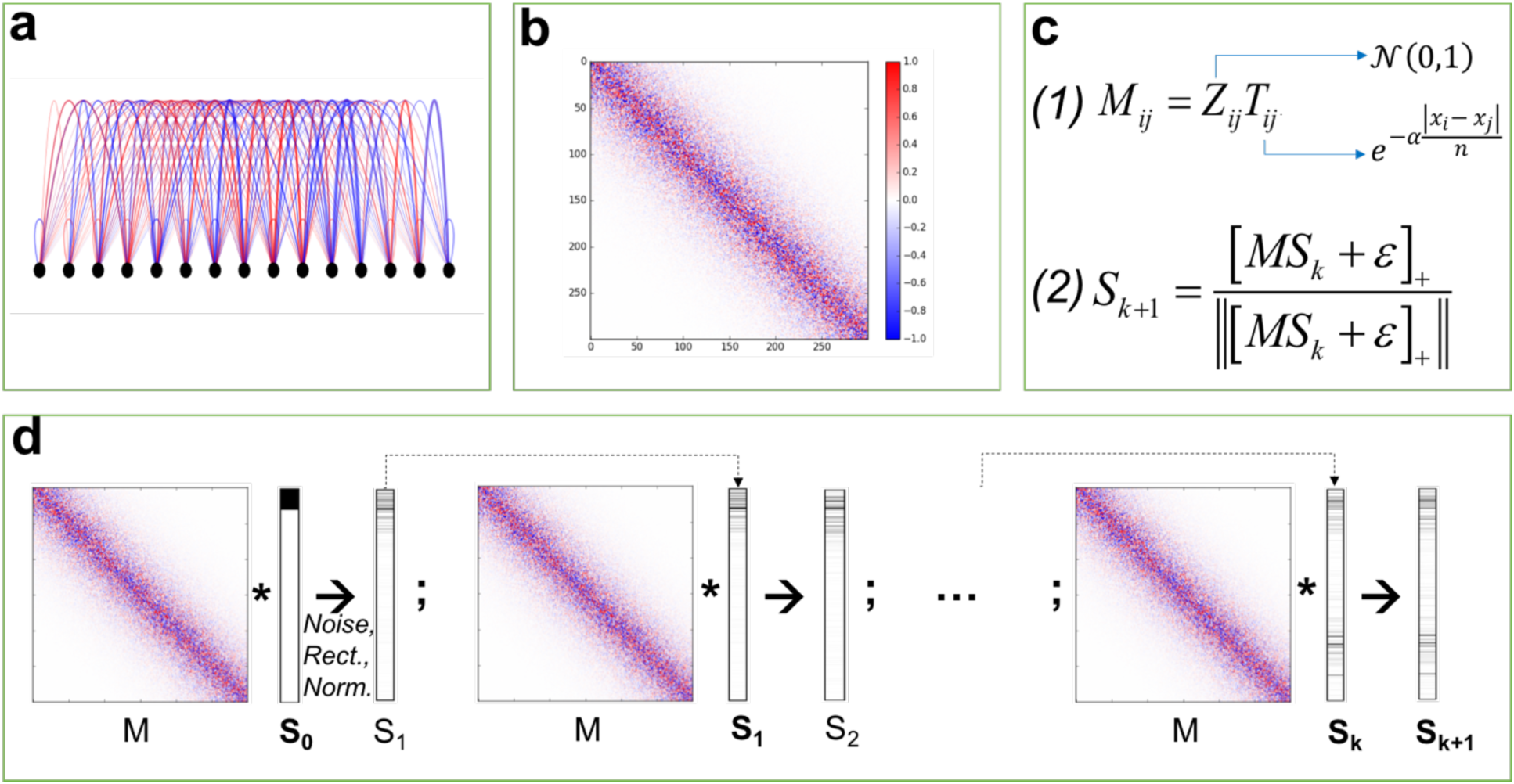
Theory of the spontaneous emergence of number sense in a random network: minimal model. **(a)** We consider a network of units regularly disposed on a segment, where each unit’s activity represents the firing of a distinct neuron. **(b-c1)** The adjacency matrix associated to the network is Gaussian (*z*_*ij*_) and local (*T*_*ij*_): the connection strength between two units is randomly drawn but scaled down exponentially as a function of their distance on the segment. **(c2-d)** Successive number states S_0_, S_1_, …, S_n_ are generated by the iterated application of the fixed adjacency matrix: starting from vector S_0_, the successor S_k+1_ of each vector S_k_ is obtained by multiplying it by matrix M, adding a small Gaussian noise, rectifying and normalizing.

Our abstract model does not process external stimuli. Rather, what we strive for in this article is to introduce an algorithm that produces a certain sequence of brain activity states (S_k_)_k=0..n_ appropriate to represent integers inasmuch as they are discrete and linked by a successor operation, as defined in Peano arithmetic, and to offer a close examination of the properties of those neural states. Given any state S_k_, the next number state S_k+1_ is obtained by multiplying S_k_ by M, adding a small Gaussian random noise term ε, before rectifying activations above zero and normalizing (Fig. 1d). In this manner, different numbers are assigned distinct vector codes, while the successor function is implemented by matrix multiplication. We note that in the cortex, rectification could be the result of competition among neurons or of firing thresholds at the cell level *(27)*, while normalization has been described as a canonical neural computation *(28)*. Critically, aside from the noise and the rectification enforced onto the system, this set-up is nothing else than the Power method, also known as “von Mises iteration” *(29, 30)*, a well-known algorithm that identifies the main eigenvector of a matrix by iterative matrix multiplication from an arbitrary vector state.

Figure 2 compares our simulation results to the electrophysiological data on number neurons *(7, 11)*. Although we start with a simple clustered vector and update it with a random matrix, units tuned to number spontaneously emerge: simulations show that, for every number *i*, including zero and up to 30, one can find units whose activity peaks at iteration *i* (see Supplementary Note 2). Furthermore, the average normalized activation of all model units that prefer a given number *i* tightly parallels the log-Gaussian tuning curves reported in actual neuronal recordings *(7)*. On a linear scale, tuning curves overlap and show increasing bandwidth and skew for increasing numbers. Once plotted on a log scale, however, tuning curves become symmetrical and of similar width (**D2**). These properties reflect the fact that under weak conditions on M and S_0_, the von Mises iteration converges geometrically (see Supplementary Note 9). Furthermore, if units are binned according to their preferred number (Fig. 2c), a U-shaped distribution emerges (**D3**), with more units selective to the lowest and highest tested numerosities, in agreement with the experimental data (Fig. 2d).

**Fig. 2.**
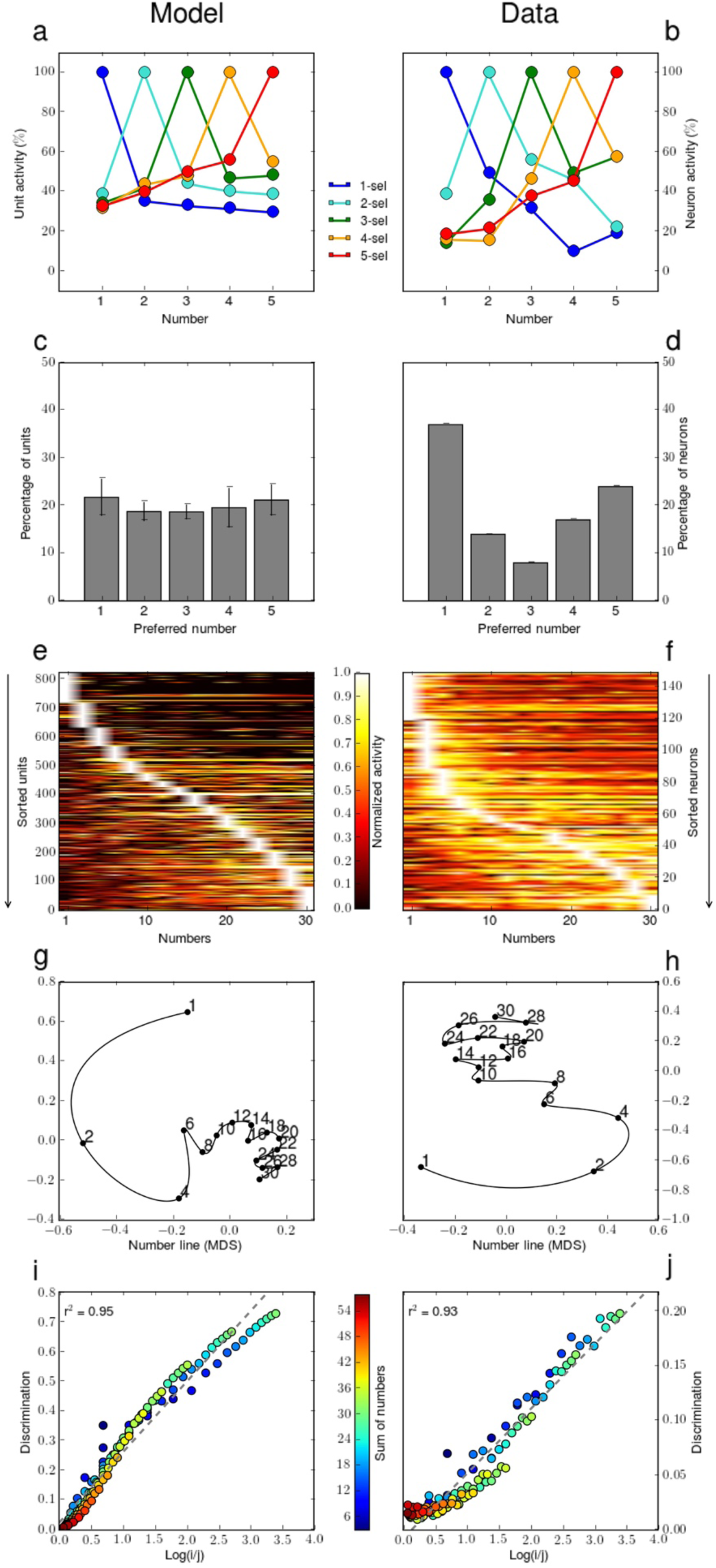
Comparison of vector states predicted by the minimal model (left column) with single-cell recordings (right column)**. (a-b)** Average tuning curves obtained by averaging the normalized activation of all units whose firing peaks to a given number (between 1 and 5). The model comprises tuned cells of increasing bandwidth and skew, similar to actual recordings [data from *(11)*]. **(c-d)** Distribution of preferred numerosity. More units are selective to the boundaries of the tested range than to mid-range numbers [data from *(11)*]. **(e-f)** Tuning to numbers 1-30 [data from *(7)*]. Units can be tuned to such large values and, when sorted by preferred numerosity, their normalized activities assume a sigmoidal distribution. **(g-h)** Reduction of neuronal state space to two dimensions using multidimensional scaling (black line = interpolation by parameterized polynomials). Number states form an oscillating and slowly converging sequence. **(i-j)** Weber-Fechner law: discriminability of number states (measured as 1-cos(S_i_,S_j_)) as a function of log number ratio. Consistent with Weber-Fechner law, the relationship is well-fitted by a linear function.

Next we ask whether the model is capable of assigning distinct vector states to larger numbers. Without changing any parameters, we produce 30 consecutive number states by successive application of matrix M to the initial vector S_0_. We then compare these theoretical vectors to the empirical vectors of firing rates of prefrontal cortex (PFC) number neurons recorded in monkeys presented with numerosities 1-30 *(7)*. Figures 2e and 2f present the normalized activities of model units and of PFC neurons, ordered by decreasing preferred number. In both cases, the sigmoidal white crests of maximal unit responses confirm the underlying U-shape distribution of number preferences previously observed for numbers 1-5. The increasing bandwidth of tuning curves is also reflected by the widening yellow regions around the white crest as numerosity increases. Interestingly, disconnected “islands” of high activity appear in some rows, suggesting that some neurons and units may be sensitive to multiple numbers.

Figure 2g and 2h show planar projections of number states in the model and in the data, obtained by multidimensional scaling. This procedure yields trajectories in state space, or multidimensional “number lines”. We observe that number lines in the data and in the model share three properties: they are compressed, with a diminishing portion of the curve devoted to increasing numbers; they are bounded, with each trajectory converging slowly to an attractor state; and they are sinuous, showing oscillations particularly for high numbers. Such oscillations are known to arise in the convergence of the von Mises iteration when the two dominant eigenvalues of the matrix are complex conjugates *(31)*.

Figures 2e and 2f also show that the Weber-Fechner law scales up to numbers 1-30: both in the model and in the data, the discriminability of number states is, to a good approximation, proportional to the log of the number ratio (**D1**). Thus, our random-matrix algorithm captures the previously reported average and large-scale properties of the number sense: the average unit represents a log-Gaussian tuned number neuron, and discriminability is determined by log-ratio.

We now describe how this minimal model accounts for additional empirical data from adult monkeys, human neonates and children of different ages.

To capture ordinal knowledge in trained monkeys *(12)*, we trained linear classifiers (SVMs) to identify the larger number amongst all 12 possible distinct pairs of vector states between 1 and 4, and then tested them on the 72 number pairs between 1 and 9, thus evaluating their capacity to generalize to numbers outside of the original training range (see Supplementary Note 5). Figure 3a shows simulated accuracies averaged over all classifiers, for each possible number pair. Matching the empirically observed performance of monkeys (Fig. 3b), the classifiers’ performance exhibits a distance effect and is above chance in comparing pairs of numbers that both fall outside of the training range 1-4 (Novel/Novel pairs, property **D4**). It may seem counterintuitive that an arbitrary vector, iteratively updated by a random matrix, produces a non-random sequence of states that contains generalizable information about numerical order. However, the presence of a spontaneous order in vector space arises naturally from the slow convergence of the “von Mises” algorithm to the main eigenvector of matrix M, and is attested by multidimensional scaling (Fig. 2g-h).

**Fig. 3.**
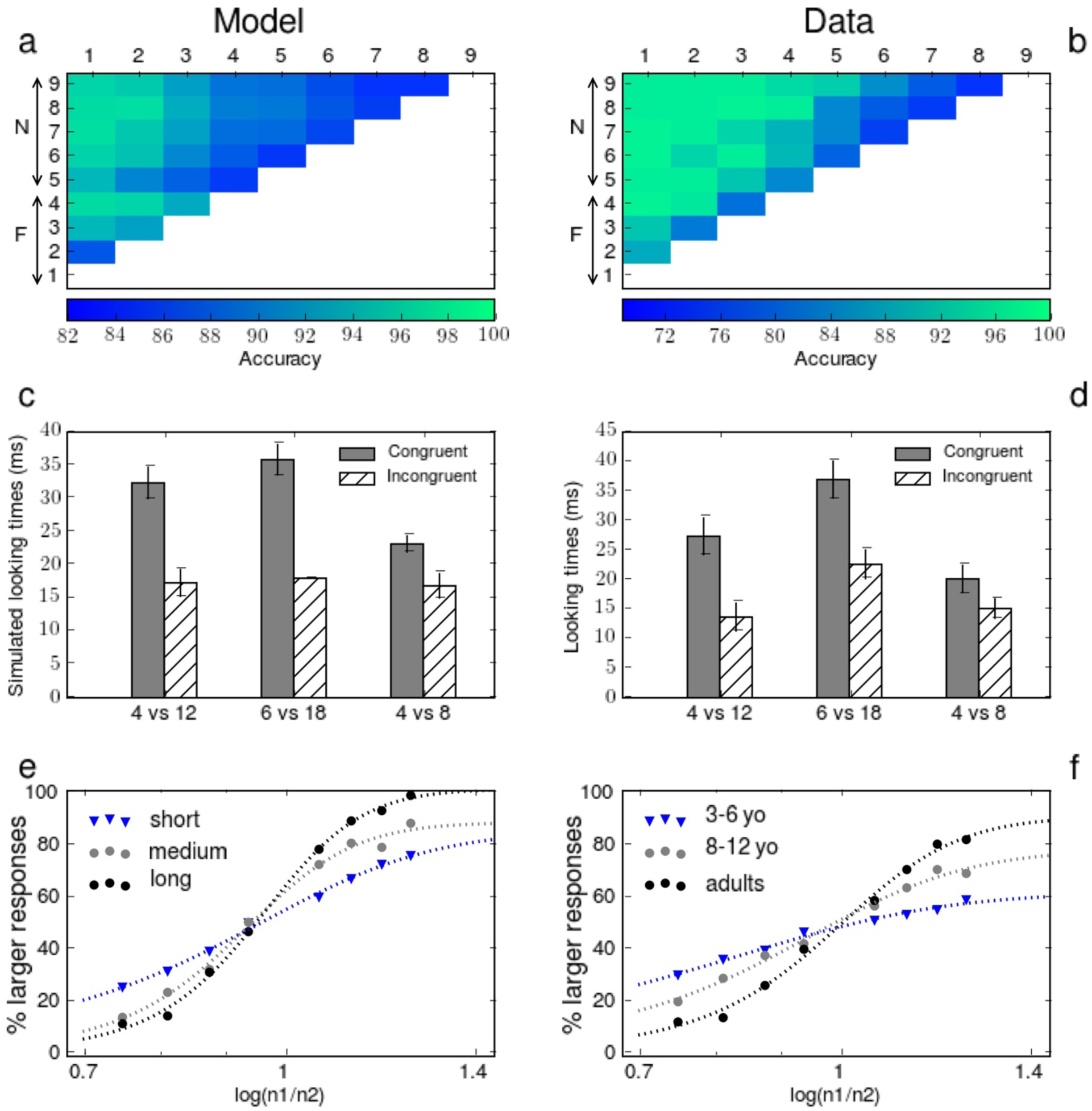
Performance of the minimal model on macroscale properties of the number sense**. (a-b)** Spontaneous generalization of ordinality judgments [data from *(12)*]. Monkeys and supervised classifiers (see Supplementary Note 6), when trained to compare numbers between 1 and 4 (marked “F” for Familiar), were able to partially generalize to a larger range of numbers between 1 and 9 (marked “N” for Novel). **(c-d)** Number discrimination in human neonates [data from *(5)*]. An unsupervised classification algorithm, trained with several noisy number states coding for the same number *n*, detects novel number states coding for another number *n*’, and its scaled performance (see Supplementary Note 5) matches the looking times of neonates in a similar numerical familiarization paradigm. **(e-f)** Improvements observed in numerosity discrimination during development are well-captured by the performance of linear classifiers at different levels of training (short: 3 runs, medium: 5 runs, or long: 10 runs), while the number states themselves remain unchanged [data from *(14)*].

Infant recovery-from-adaptation paradigms *(5)* can be simulated by training a one-class support vector machine on noisy vector states obtained from different runs of the same model for the same numerosity (see Supplementary Note 6). When tested with a new vector state that either matches or differs from the familiar numerosity, emulating the procedure in *(5)*, the classifier rejects novel numerosities more than habitual ones, and this effect reproduces infant looking times (Fig. 3c). Critically, this response to numerical novelty is larger when the habituation and deviant numerosities differ by a 3:1 ratio than when they differ by a 2:1 ratio, consistent with Weber’s law and in agreement with empirical data in neonates (Fig. 3d, **D1, D5**).

Using the same logic, we simulated the improved number discrimination that accompanies human development and education (**D6**) *(14, 15)* (see Supplementary Note 7). According to our theory, vector representations for numbers are innate and stable. Why, then, does the behavioural Weber fraction decrease with age and education? We attribute this evolution to a progressive refinement in the way numerical information is extracted from the neural population activity. This postulate fits with direct electrophysiological evidence that parietal number neurons are already present in untrained monkeys, in proportions that remain unchanged by training, and that only the proportion of prefrontal number neurons is enhanced when monkeys are trained in a numerical match-to-sample task *(32)*. We therefore simulated the development of numerical comparison abilities as the training of a classifier to decide which of two numbers is larger. When performance is assessed early on during training, classifiers initially display poor discrimination, with performance varying as a shallow function of the log ratio of the numbers involved, mimicking young children’s data. With additional training, discrimination is increasingly better fitted by a steep sigmoidal, reflecting sharper number judgements and a shift in the apparent Weber fraction (Fig. 3e-f, **D6**).

We note that, if the same population of neurons also contained overlapping codes for non-numerical parameters such as physical size *(33-35)*, our hypothesis that development consists in a progressive focusing of decision-making on the relevant numerical dimension could provide a natural explanation for why children initially confuse number with other physical dimensions (e.g. Piaget’s classical “number-conversation errors”), and why such non-numerical interference decreases in the course of development *(36)*. We did not explicitly simulate this aspect of our theory here, however, because this would require additional assumption about the coding of those non-numerical dimensions.

### The extended number sense model

The above model is deliberately abstract and lives in a space of dimension 1. In a second implementation of our theory, we aimed to study whether the same properties would hold in a more realistic model of neuronal dynamics, including a network of excitatory and inhibitory units embedded within a cortical plane, with sparse and local connectivity (Fig. 4). This extended model follows Dale’s principle *(37)*: units can either be excitatory or inhibitory, but not both (see *ref. 38* for a similar application of Dale’s principle to random networks). As observed in the number system, excitatory units in the extended model outnumber inhibitory units 4 to 1 *(2)*. The absolute value for each entry M_ij_ is still given by a Gaussian random variable whose amplitude drops exponentially with the distance between units. We also introduce a more realistic sparsity constraint on all network connections. We now show that this model retains all the properties of the minimal model, and also accounts for several additional phenomena.

**Fig. 4.**
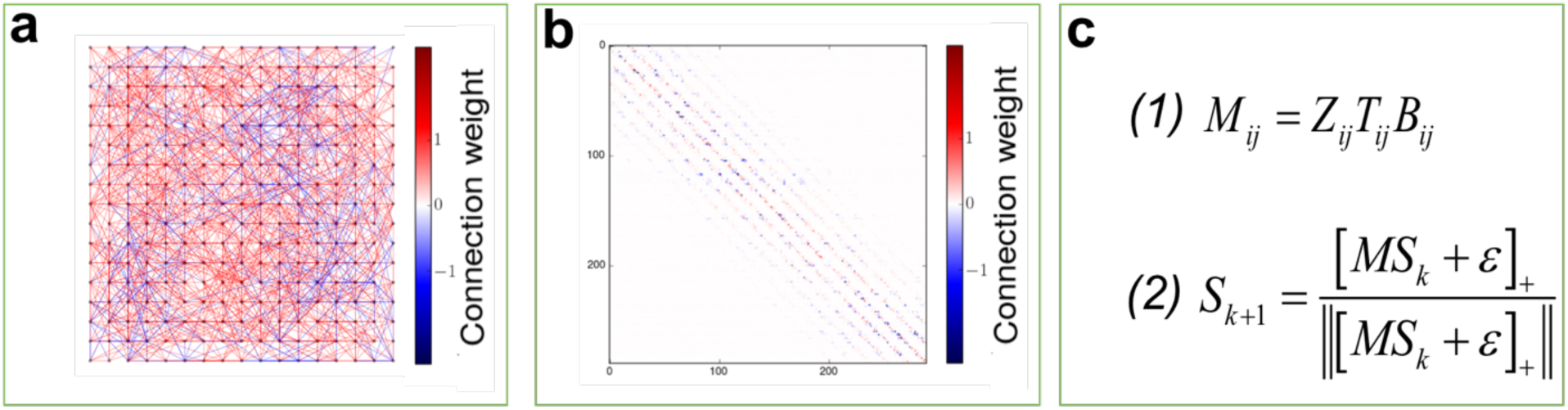
Theory of the spontaneous emergence of number sense in a random network: extended model. **(a)** We consider a 2D grid of excitatory and inhibitory units, sparsely connected (strongest connections shown, excluding self-connections). **(b-c1)** The adjacency matrix associated to the network is Gaussian (*z*_*ij*_), sparse (*B*_*ij*_) and local (*T*_*ij*_): the connection strength between two units scales down exponentially with their distance on the grid. A unit can either be excitatory (red matrix lines) or inhibitory (blue matrix lines) but not both. **(c2)** The dynamics of the model is otherwise unchanged with respect to the minimal model.

Figure 5 assesses the behaviour of the extended model on the same properties as Fig. 2. It shows that number states in the extended model continue to display the same properties as those of number neurons. Specifically, the tuning curves overlap and exhibit skews that increase with numerosity (**D2**, Fig. 5a), more units are selective for the first and last number in the tested range (**D3**, Fig. 5c, 5e), number states still appear to converge exponentially to an attractor (Fig. 5g), and the Weber-Fechner law is conserved (**D1**, Fig. 5i). In addition, the distinction that we now make between excitatory and inhibitory units captures more detailed properties of number neurons. Figure 5k shows that model responses are similar for nearby pairs of excitatory units, but complementary for pairs of excitatory and inhibitory units (see Supplementary Note 4). This interaction closely mirrors the empirical relationship of tuning curves for adjacent number neurons in the PFC (Fig. 5l), as reported for broad and narrow spiking cells (corresponding to putative excitatory and inhibitory cells) recorded from the same electrode *(2, 39)*. This finding suggests that even detailed properties of the micro-circuitry of the approximate number system can be captured, in first approximation, by our simple model.

**Fig. 5.**
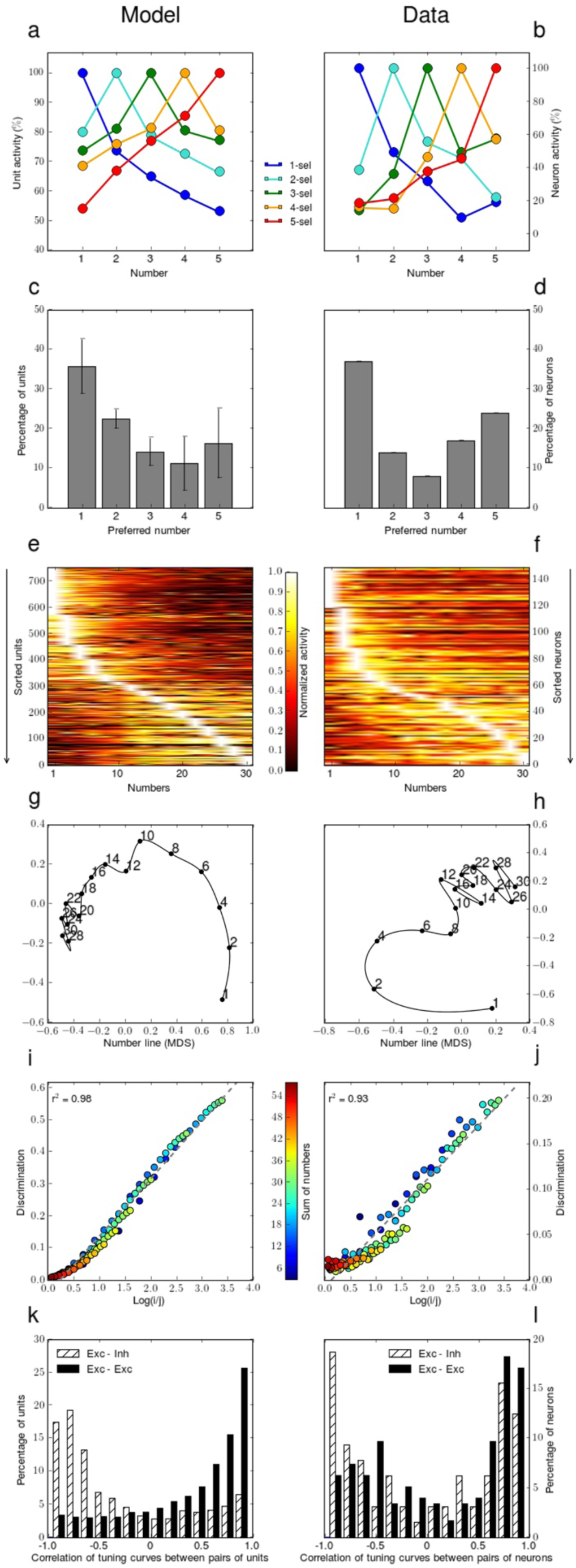
Comparison of vector states predicted by the extended model (left column) with single-cell recordings (right column). The model continues to account for the same single-cell properties of the number sense as previously. In addition, the histograms of tuning curves similarity between neighbour units in the model (**k**) show similar distributions to those reported for adjacent neurons (**l**) [data from *(34)*].

Figure 6 likewise evaluates the extended model on the same number sense properties as Fig. 3, and shows that it can still account for properties **D4** (Fig. 6a), **D5** (Fig. 6c), and **D6** (Fig. 6e). Importantly, our 2D model now enables the investigation of mesoscale aspects of the number sense as detected by high-field imaging in human adults. We thus ask whether the kind of spontaneous ordering of number states shown by the model could account for the emergence of “numerotopy” *(16)*, i.e. the similar location of voxels selective for similar numerosities (**D7**). When the initial vector S_0_ is clustered on the left side of the grid, the preferred number diffuses locally from that initial cluster (Fig. 6e), mirroring the medial-to-lateral gradient of selectivity observed using fMRI in human subjects (Fig. 6f). On average, the location of a unit along the medial-to-lateral axis correlates with its number preference (**D7,** r_pearson_=0.95, p_pearson_<<0.001; excluding zero selective units: r_pearson_=0.86, p_pearson_<<0.001), as observed experimentally (Fig. 6g-h). According to our model however, such a medial-to-lateral direction is not universal but contingent upon the location of the units which are activated in the initial state S_0_: different locations would produce different gradients, depending on the diffusion opportunities offered by the source. In fact, an initial vector S_0_, containing multiple randomly placed clusters of units selective to zero, results in multiple coexisting gradients that run in opposite directions on different parts of the simulated cortex, as is also manifest in the data *(16)* (Fig. 6i-j). Similarly, our simulations show that the convexity or concavity of the gradient is not universal, but depends on the spread of the zero-selective cluster. Overall, the diffusion process by which we explain numerotopy is not unrelated to the self-organizing Turing reaction-diffusion model *(40)*, although in our case the large-scale topography of number regions only obtains after smoothing of the selectivity map by a Gaussian filter. We thus predict that higher-resolution imaging should uncover a more intertwined topography of number preferences (see Supplementary Note 8). The fact that voxel-level properties such as numerotopy can be accounted for by exactly the same model as single-cell effects would also suggest that the approximate number system is self-similar across scales.

**Fig. 6.**
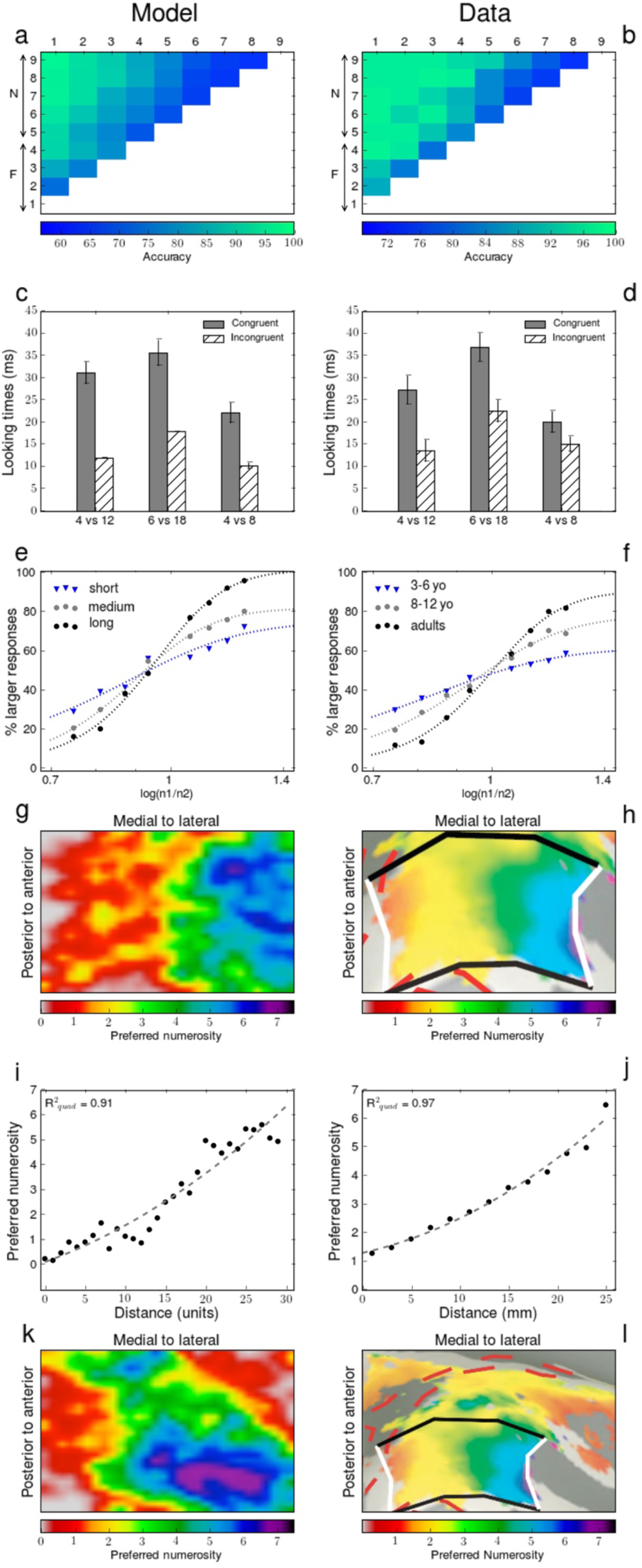
Performance of the extended model on macroscale properties of the number sense. The extended model still accounts for spontaneous generalization of ordinality judgments **(a-b)** [data from *(12)*], for number discrimination in human neonates **(c-d)** [data from *(5)*], as well as for the improvements in numerosity discrimination reported during development **(e-f)** [data from *(14)*]. In addition, the model explains numerotopy in human adults **(g-h)** [data from *(15)*]. Because vector states are considered as two-dimensional patterns on a cortical grid, the iteration algorithm implements a stochastic diffusion process. (**i-j**) Since the initial state S_0_ is clustered at one end, preferred numerosity increases monotonically with distance on the cortical surface, as observed experimentally in human parietal cortex. (**k-l**) When two diagonally opposed activation sources exist in the initial state, the model produces two selectivity gradients, which run in opposite directions on the bottom and top parts of the grid, but “coalesce” in the centre to form a large region of intermediate selectivity (here numbers 2 and 3).

### Drawing quantitative predictions from the theory: Two critical tests

We now offer two quantitative predictions that set our theory distinctly apart from other alternatives. First, our theory predicts that individual units can respond to more than one numerosity (Fig. 2e), suggesting that the log-normal model more often captures an average firing rate profile over many neurons with the same preferred numerosity. To test this prediction and quantify the proportion of “multipeak” tuning curves in real data, we conducted a post-hoc analysis on all neurons previously identified as number selective in the sample period of ref. *(7)* (see Supplementary Note 3). Given a neuron with preferred numerosity *n*, the general logic of this analysis was to grant another peak at numerosity *n’* whenever the firing rate of the cell did not differ significantly in response to *n* and *n’*, but was significantly reduced for numerosities between *n* and *n’*.

We discovered that 10.6% of number neurons exhibit unambiguous tuning to multiple numbers, against 4.6% units in the minimal model and 3.4% in the extended model (see examples in Fig. 7). We note that units with multiple number preferences are a generic behaviour of our random-matrix theory: they appear even in the absence of noise, as a counterpart at the unit level of the oscillatory behaviour previously reported at the population level. The fact that actual number neurons conform to this prediction thus provides strong support for the random-matrix theory.

**Fig. 7.**
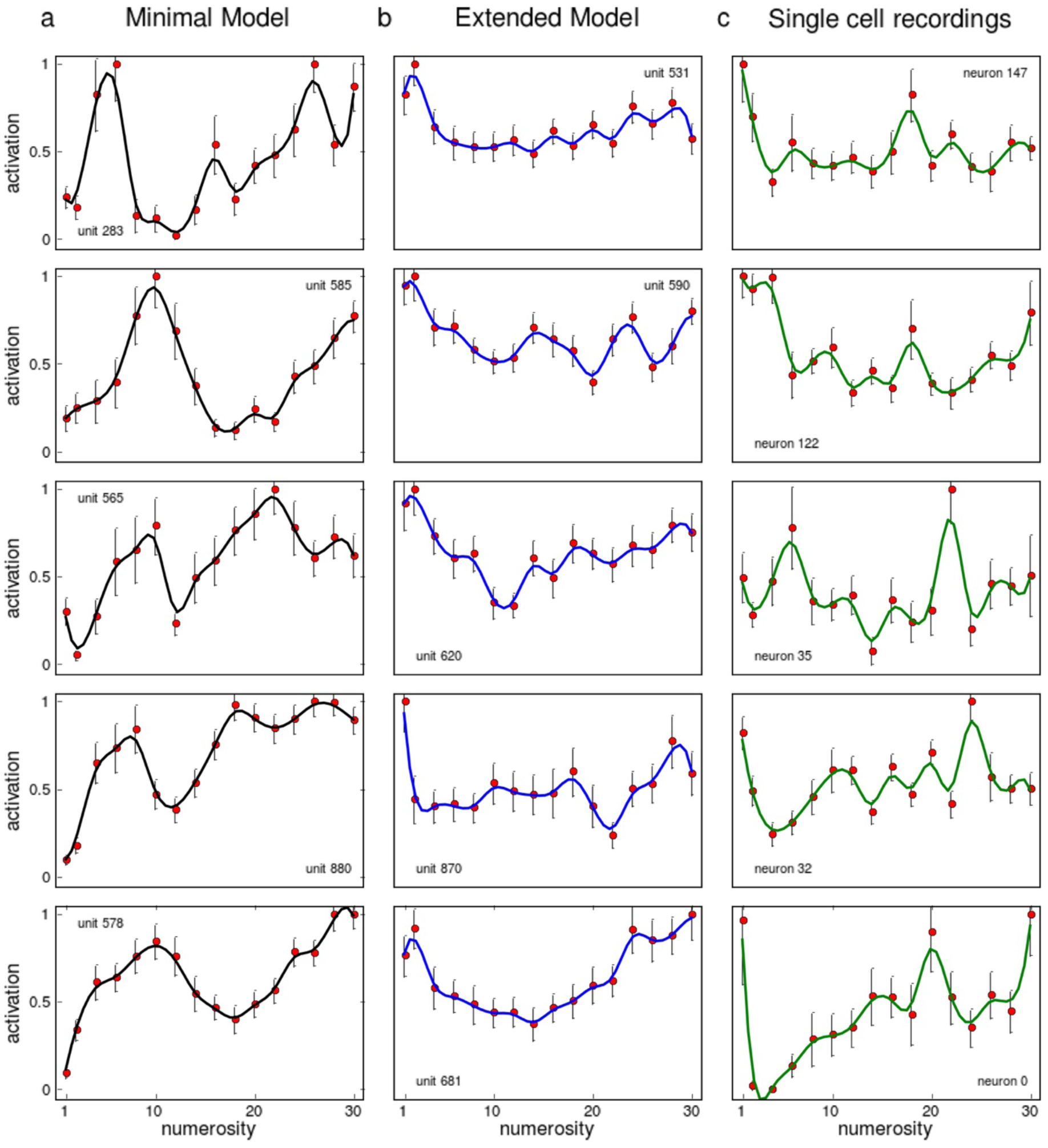
Examples of tuning curves indicating the presence of “multipeak” responses in the models and in actual recordings -data from *(7)*. **(a)** Our conservative post-hoc analysis shows that 4.56% of units in the minimal model exhibit unambiguous tuning to multiple numbers, and **(b)** 3.44% in the extended model. **(c)** The same analysis performed on the single cell recordings from *(7)* reveals 10.60% of multipeak number neurons. Mean firing rates/activities across the sample period (red dots) were smoothed by a Gaussian kernel and interpolated (solid curves). Error bars show the standard error of the mean.

Our second quantitative prediction concerns the fundamental assumption of our theory, that of a random band matrix which implements a successor function. Because this matrix links number codes together, it is legitimate to ask whether its dimension n and bandwidth W would impact on number sense. In fact, a long standing conjecture *(41)* in random matrix theory states that for random band matrices whose underlying space is of dimension 1, a critical bandwidth W* exists that marks a sharp phase transition between two regimes: a strongly disordered, “insulator” regime for bands smaller than 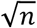, where eigenvectors are localized in the sense that most of the norm of the vector is carried by a few elements, and a weakly disordered, “metallic” regime for bands larger than 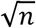, where eigenvectors are delocalized (i.e. more distributed). Though there is no exact counterpart of this conjecture for the more biologically motivated matrix used in the extended model, some results for dimension 2 random band matrices point to a critical bandwidth that would scale like 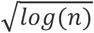 *(42)*.

We therefore examined whether the models changed behaviour when W approaches the critical bandwidth, and discovered that this is precisely the point where Weber-Fechner’s law holds with the greatest precision. Figure 8 shows how Weber-Fechner scores vary as a function of n and W from trials to trials. Both models exhibit a smooth landscape of Weber-Fechner scores. In the minimal model, global maxima (white crosses) are situated strikingly close to the conjectured critical bandwidths of 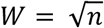, (Fig. 8a, dashed curve). In the extended model, there is a sharp transition from high to low scores after a critical bandwidth that also scales approximately like the square-root of n. Note that in the extended model the precise shape of the relation between critical W and n depends exquisitely on the structure of the initial zero state (see Supplementary note 11), though a common feature is that of critical bandwidths increasing monotonically with n. Thus in our theory the Weber-Fechner law appears to be linked to conjectures on phase transitions in random band matrices.

**Fig. 8.**
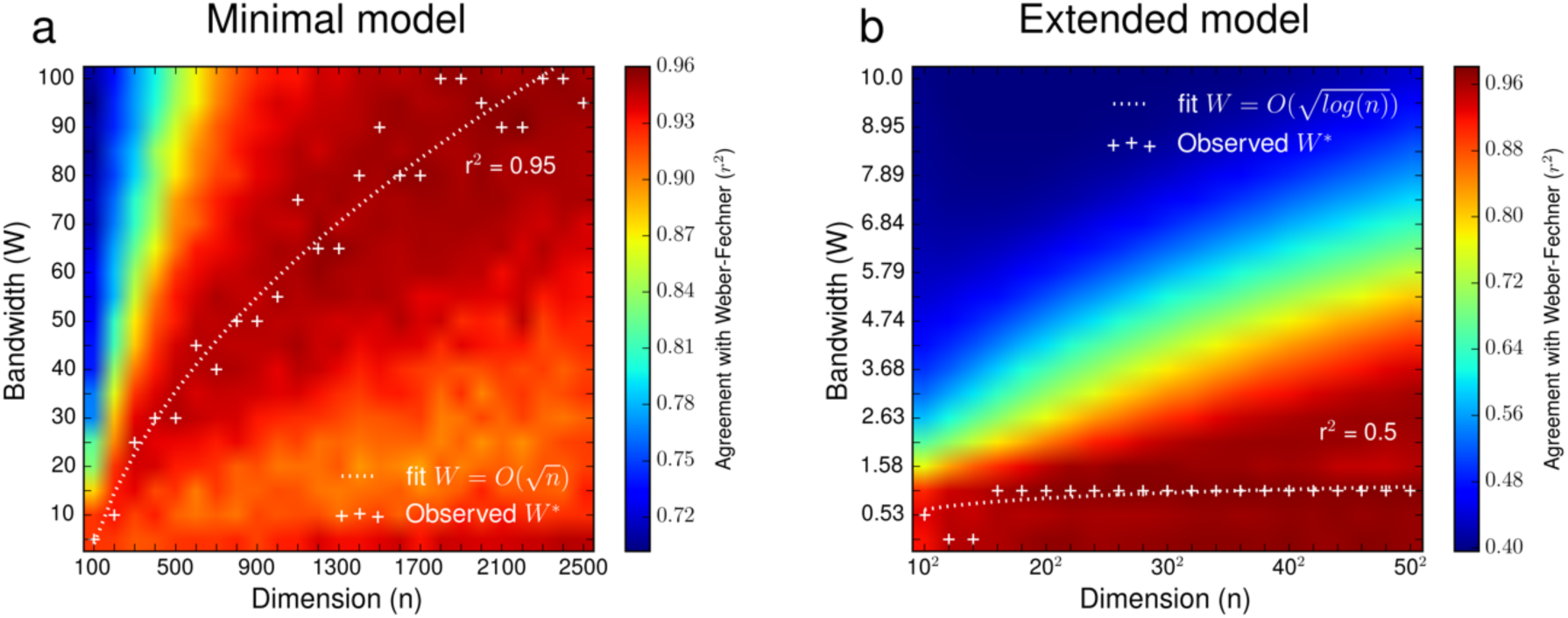
Predicted phase diagrams for Weber-Fechner scores as a function of dimension n and bandwidth W. **(a)** Weber-Fechner scores are generally high in the minimal model, but optimal bandwidths are well-fitted by a curve proportional to 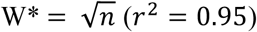, which is conjectured to mark a phase transition for 1D random band matrices *(41)*. **(b)** In the extended model, there is a sharp transition from high to weak scores past a critical border that also scales like the square root of n (notice the quadratic x-axis), although the optimal bandwidth is not very well captured by the relation 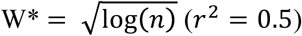, expected from the most applicable conjecture for a phase transition in 2D random band matrices *(42)*.

This prediction sets our theory distinctly apart from those which hold that number representations do not exist in the abstract, but are transient constructs computed from combinations of low-level stimulus properties (e.g., in the case of visual dot displays, total surface area or density, ref. 18): according to these views one would not expect any simple relation to hold between the composite representations obtained for all consecutive numbers, let alone any link to criticality phenomena in random band matrices. Testing this prediction would require to estimate the properties of matrix M, for instance by computing the cross-correlation matrix between a significant fraction of number neurons as a function of their cortical distance, for different subjects, trials and numbers. High density 2D arrays of multielectrodes could be sufficient for the task, as they are known to provide sufficient resolution to distinguish between inhibitory and excitatory cells (*26*), which would be necessary to test the predictions of the extended model.

## Discussion

We have shown how a simple theory, based on random matrices, reproduces all essential aspects of number sense, including those for which learning cannot be assumed. The theory conserves its explanatory power across a range of parameters and implementation choices (see Supplementary Note 9), but its key ingredients lie in the assumptions that (i) each number is coded by a sparse, rectified and normalized vector; (ii) the vectors for consecutive numbers are iteratively linked through multiplication by a fixed random band matrix M. Not only does this theory provide a better account of number neurons than our previous log-Gaussian approach, particularly in predicting and accounting for the new discovery of multipeak number neurons, but it also endows number codes with a “matrix” successor function (+1 is implemented by one application of M). In the present view, distinct numbers are not simply indexed by distinct populations of number neurons, but these neurons actually reflect a vector-based system of interrelated symbols linked by a fixed successor operation *(43)*. Indeed, other basic arithmetic operations such as +2 or −1 could be implemented by simple matrix operations (M^2^, M^−1^). Our theory therefore offers a first glimpse of how uneducated adults, monkeys and even young infants, without training, could be endowed with a sensitivity for non-symbolic arithmetic and its violations *(44-46).*

Our neural implementation of Peano’s successor function, a fundamental requisite of arithmetic, should not be confused with the successor function familiar to developmental psychologists, i.e. the laborious acquisition, during childhood, of a system of number symbols one, two, three… linked through counting. While our model accounts for how human and non-human primates, as well as other species, can be innately endowed with approximate number vectors, much remains to be understood about how these vectors are linked, on the perceptual side, to sets of objects, and on the abstract side, to symbols such as Arabic numerals or verbal number words.

Although our theory for the origins of number vectors includes realistic properties such as Dale’s principle, sparsity of neural activity patterns, and sparse and distance-dependent connectivity, it describes brain activity at an abstract mathematical level similar in spirit to the classical Hopfield model for neural networks *(47)*. This abstract nature need not be a disadvantage, for it also constructs a bridge between the mathematical field of random matrices and the neuroscience of number. Specifically, our simulations suggest that the Weber-Fechner law for numbers arises from a critical network connectivity in an eigenvector algorithm that runs on a population of neurons (see Fig. 8 and Supplementary Note 10).

Among the many steps that one might take towards more realistic models, using spiking units and differentiating the firing rate levels and connectivity ranges for excitatory and inhibitory units would no doubt help capture the micro-circuitry of number neurons. Most importantly, the present proposal leaves unspecified the nature of the gating mechanism that would implement the required step-like application of matrix M each time one moves from numerosity *i* to *i+1*. This assumption sets our theory apart from otherwise similar proposals of “liquid computing” or “echo-state networks” in which an internal vector representation is allowed to continuously evolve under the continuous application of a fixed and possibly random dynamics *(48-50)*, and which capture several aspects of the neural representation of time *(51)*.

One challenge facing our theory is to reconcile its inherent sequentiality with the absence of observed delays in neural firing patterns for different numerosities *(4, 11)*. By analogy with the dynamic waves of spontaneous retinal activity that ultimately generate static retinotopic visual maps *(52)*, we suggest that the postulated matrix iterations may occur dynamically during gestation, shaping number sense by generating ordered vector states that are only later being associated in parallel with external stimuli. A more detailed description of this association would require specifying how our abstract model interacts with the computational machinery that processes actual stimuli in the visual and auditory pathways. While the organization of such perceptual processing has been the main focus of previous theoretical models of number sense *(17-20)*, the main virtue of the present theory is to address the problem from the opposite direction: we demonstrate how structured neural codes for number can emerge from sheer randomness through an endogenous mechanism that is minimal enough to be specified genetically. Such a spontaneous emergence, without training or specific sensory interactions, fits with the presence of number neurons in untrained animals *(3-4)*, of parietal-lobe responses to number in human infants *(53)*, and of normal parietal brain networks for number sense even in congenitally blind subjects *(54)*.

The theory makes several novel predictions. The first, which we verified here for the first time through a reanalysis of existing data, is that individual number neurons can exhibit multiple peaks of numerosity preference (Fig. 7). It mirrors the recent finding of multipeak time-coding cells in the hippocampus (*55*), and lends support to the notion that similar iterative processes could be at work in the cognition of time and space (*51*). A second prediction is that the cross-correlogram between number neurons should be stable across numbers, and in normal subjects should assume the form of a random band matrix with bandwidth W set close to criticality, and away from criticality in subjects with an impaired number sense (Fig. 8). Third, numerotopy should only be an average property, and imaging with higher fields should reveal distributed number tuning in parietal neurons, as well as complex multipeak neurons. Forth, number neurons and numerotopy should already be present in the newborn brain. Given the presence of number sense in many species, our theory hints that the nervous system may have been harnessing the properties of random matrices as a self-organizing representational system for millions of years.

## Methods

Our random-matrix theory of the number sense relies on an iterative process that constructs a sequence of number states of dimension n, using a (n × n) random band matrix M as a successor function, and a clustered initial state.

### The minimal model

In the minimal model, each unit *i* = 1, …, *n* is disposed regularly on a line, and the distance between two units is defined as their absolute distance. Matrix M constitutes the adjacency matrix of the network defined on this line: each entry *M*_*ij*_ is a Gaussian random variable, rescaled so as to decrease exponentially with the distance between units i and j:

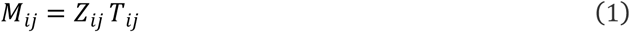

Where

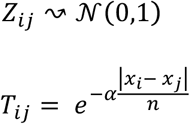

The minimal model has very little structure: in absence of locality (***α*** = 0), we recover a standard Gaussian matrix.

Starting from a deliberately simple initial state S_0_, in which activation is confined to the leftmost part of the line, each new state is obtained by multiplying the current one by M, adding a small standard Gaussian noise ε with amplitude ***α*_*ε*_**, rectifying, and normalizing:

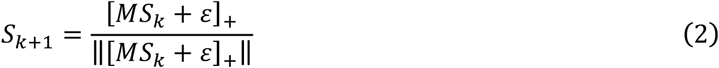

Where [.]_+_ stands for rectification above zero.

Hence, the minimal model has 3 main parameters: ***n, α, α*_*ε*_**. In addition, the initial state requires another parameter ***r*_*0*_**: before normalizing, all units in the initial state are set to 0, except for the leftmost ***r*_*0*_ *n*** units which are set to 1. Table 1 gives an exhaustive list of parameters in the minimal model.

**Table 1.** Parameters of the minimal model.

| *Parameters* | *Values* | *Description* | *Default* |
| --- | --- | --- | --- |
| $\mathbf{n}$ | $\mathbb{N}$ | Dimension of vector state | 900 |
| $\alpha$ | $\mathbb{R}^+$ | Locality of adjacency matrix | 30 |
| $\mathbf{a}_\epsilon$ | $[0, 1]$ | Amplitude of Gaussian noise in vector state | 0.01 |
| $\mathbf{r}_0$ | $[0, 1]$ | Proportion of initially active leftmost units | 0.1 |

### The extended model

In the extended model, each unit *i* = 1, …, *n* is embedded in a 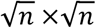 square grid, with integral coordinates *(x*_*i*_, *y*_*j*_) given by 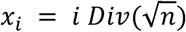 and 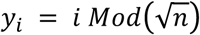. The distance between two units is defined as the L2 norm. The matrix M now constitutes the adjacency matrix of the network defined on this grid. Each entry *M*_*ij*_ is a half-normal random variable, rescaled so as to decrease exponentially with the distance between units i and j, and multiplied by a Bernoulli variable:

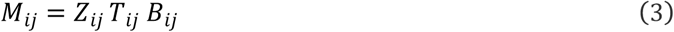

Where

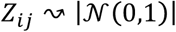

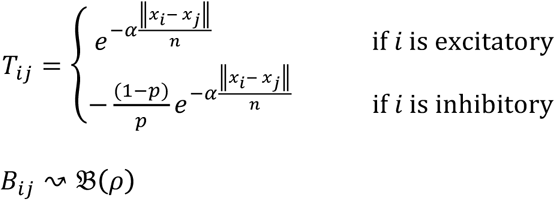

Parameter *p* controls the proportion of inhibitory units, while parameters ***ρ*** and ***α*** respectively control the density and locality of the system. Notice that inhibitory weights are rescaled so as to ensure that the adjacency matrix always has mean zero, despite the imbalance between excitatory and inhibitory connections (when *p* ≠ 0.5). The dynamics of the extended model are unchanged, and new model states are still iteratively obtained by application of equation (2).

The initial state in the extended model is still clustered and stable across trials, but the activation pattern on initially active units is now modelled as a Gaussian 2D bump of activation whose centre varies uniformly across subjects between [0, *nr*_0_] on the x-axis, and [0, *n*] on the y-axis. The extended model therefore has 6 parameters in total**: *n, p, ρ, α, α*_*ε*_**, and ***r*_*0*_,** listed in Table 2.

**Table 2.**
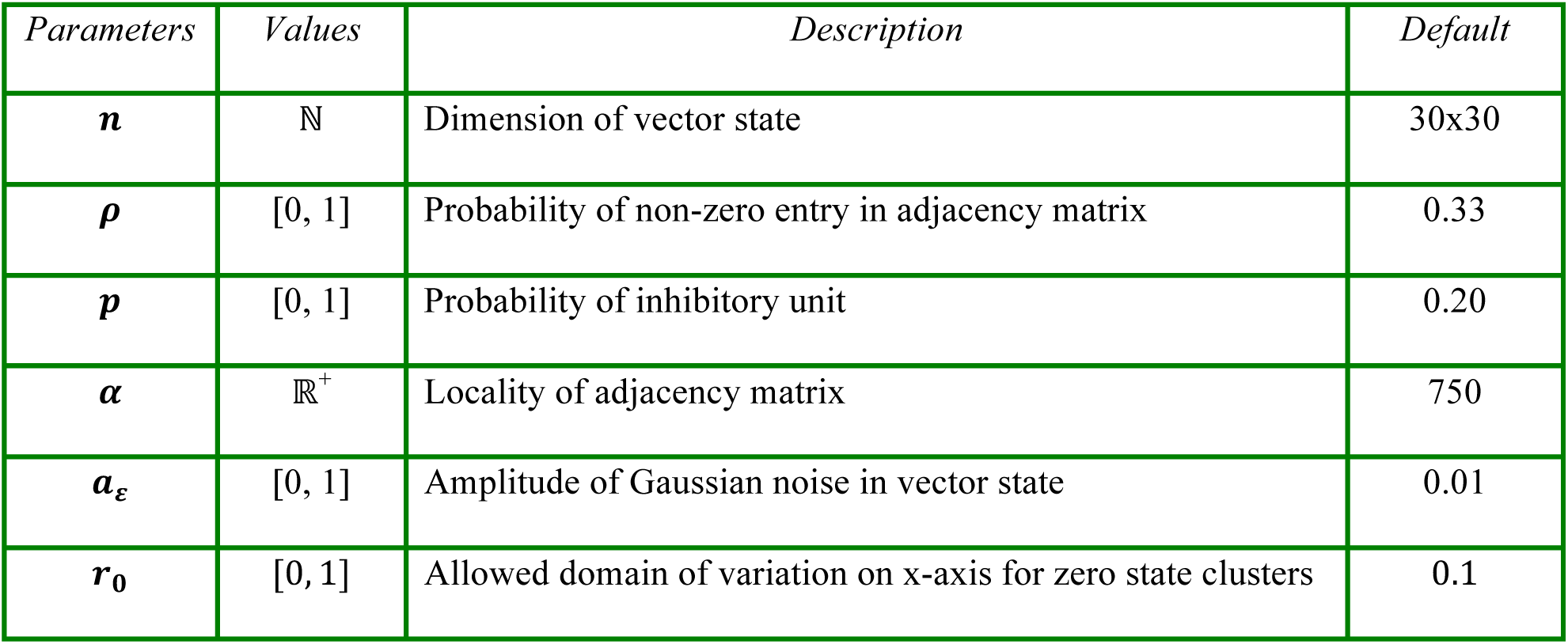
List of parameters in the extended model.

### General simulation procedure

In all of our simulations, we distinguish between different subjects (captured by model runs with different initializations of the random matrix M and different initial states), different trials (model runs with the same matrix and same initial state) and numerosities (model runs with the same M and the same zero state, for a number of iterations). The exact numbers of subjects, trials and numerosities is adapted to the experiment under simulation. Unless otherwise specified, all simulations were run with the default values given in Tables 1 and 2.

### General simulation procedure

In all of our simulations, we distinguish between different subjects (captured by model runs with different initializations of the random matrix *M* and different initial states), different trials (model runs with the same matrix and same initial state) and numerosities (model runs with the same M and the same zero state, for a number of iterations). The exact numbers of subjects, trials and numerosities is adapted to the experiment under simulation. Unless otherwise specified, all simulations were run with the default values given in Tables 1 and 2.

### Code availability

The full python code used in the simulations is available upon request to the first author, and will be made publicly available on the Unicog website (http://www.unicog.org/biblio/) after publication of the article.

## Acknowledgments

We gratefully acknowledge V. Izard, J. Touboul, E. Eger and G. Dubach for helpful discussions, V. Izard, E. Brannon and B. Harvey for data sharing. This work was supported by INSERM, CEA, and the Bettencourt-Schueller Foundation. TH was supported by the European Commission Seventh Programme (FP7/2007-2013) under grant agreement 604102 (Human Brain Project).

